# The WAVE complex forms linear arrays at negative membrane curvature to instruct lamellipodia formation

**DOI:** 10.1101/2024.07.08.600855

**Authors:** Muziyue Wu, Paul Marchando, Kirstin Meyer, Ziqi Tang, Derek N. Woolfson, Orion D. Weiner

## Abstract

Cells generate a wide range of actin-based membrane protrusions for various cell behaviors. These protrusions are organized by different actin nucleation promoting factors. For example, N-WASP controls finger-like filopodia, whereas the WAVE complex controls sheet-like lamellipodia. These different membrane morphologies likely reflect different patterns of nucleator self-organization. N-WASP phase separation has been successfully studied through biochemical reconstitutions, but how the WAVE complex self-organizes to instruct lamellipodia is unknown. Because WAVE complex self-organization has proven refractory to cell-free studies, we leverage *in vivo* biochemical approaches to investigate WAVE complex organization within its native cellular context. With single molecule tracking and molecular counting, we show that the WAVE complex forms highly regular multilayered linear arrays at the plasma membrane that are reminiscent of a microtubule-like organization. Similar to the organization of microtubule protofilaments in a curved array, membrane curvature is both necessary and sufficient for formation of these WAVE complex linear arrays, though actin polymerization is not. This dependency on negative membrane curvature could explain both the templating of lamellipodia and their emergent behaviors, including barrier avoidance. Our data uncover the key biophysical properties of mesoscale WAVE complex patterning and highlight an integral relationship between NPF self-organization and cell morphogenesis.

## Introduction

Plasma membrane protrusions are crucial for many aspects of cell physiology. For example, endothelial cells use finger-like membrane protrusions called filopodia to probe their environment and facilitate cell-cell/cell-substrate adhesion (Khurana & George, 2011; Sagar et al., 2015; Ventura et al., 2022), while migratory cells build flat sheet-like membrane protrusions called lamellipodia to power cell motility (Innocenti, 2018; Krause & Gautreau, 2014). Actin rearrangements play a key role in the spatial and temporal regulation of these membrane deformations. Nucleation promoting factors (NPFs) help specify when and where actin is polymerized (Chesarone & Goode, 2009; Pollard & Borisy, 2003; Rottner et al., 2017; Takenawa & Suetsugu, 2007). The neural Wiskott-Aldrich syndrome protein (N-WASP) orchestrates unbranched actin bundles that support spiky and narrow filopodial structures (Miki et al., 1998; Nemethova et al., 2008), while the WASP family verprolin homologous protein (WAVE) complex orchestrates sheet-like branched actin networks that underlie broad and thin lamellipodial structures (Fritz-Laylin et al., 2017; Leithner et al., 2016; Machesky & Insall, 1998; Machesky et al., 1999; Steffen et al., 2004; Weiner et al., 2007).

How do different NPFs instruct different patterns of membrane protrusion? Biochemical reconstitutions using different architectures of micropatterned surfaces coated with the same NPF have provided some insights into the relation between actin network structures and NPF organization patterns. While a dot-like NPF organization drives finger-like actin networks, a linear pattern of the same NPF drives lamellipodial-like actin networks *in vitro* (Boujemaa-Paterski et al., 2017; Carlier et al., 2003). This suggests that the spatial organization of NPFs could dictate the resulting morphology of actin assembly and cell protrusion (**Figure 1**).

**Figure 1.**
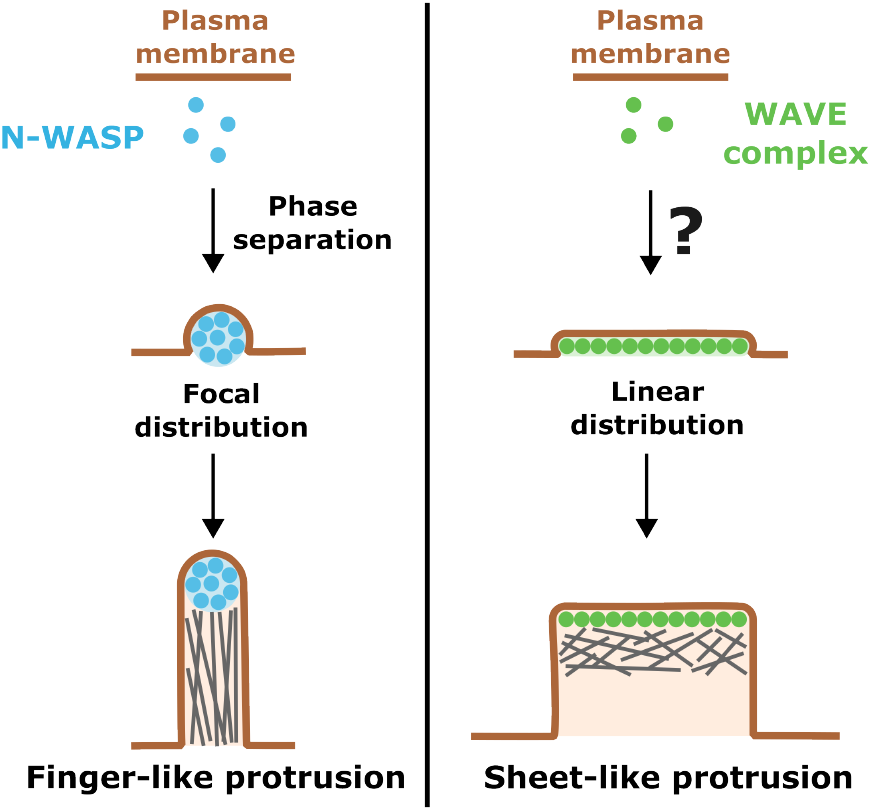
How does the WAVE complex achieve the linear distribution needed for lamellipodia formation? Different nucleation promoting factors (NPFs) form different distribution patterns on cell membranes to instruct their characteristic membrane protrusion structures. Left: N-WASP (blue) forms focal distributions by undergoing phase separation with binding partners on cell membranes (Banjade & Rosen, 2014), providing the proper distribution for building finger-like membrane protrusions. Right: The WAVE complex (green) builds sheet-like lamellipodial structures, but how it achieves the needed linear organization is not well understood.

For the *in vitro* studies, NPF distribution is typically externally constrained (Boujemaa-Paterski et al., 2017; Carlier et al., 2003). For living cells, NPF distribution is internally specificized through self-organization. For example, N-WASP proteins phase separate to form focal structures on the plasma membrane that direct the extension of filopodia (Banjade & Rosen, 2014; Case et al., 2019; Ward et al., 2004). In contrast, to build a flat sheet-like lamellipodia protrusion, the WAVE complex needs to assemble into a linear pattern on the plasma membrane (**Figure 1**). The multivalent interactions that generate N-WASP phase transition are relatively well-understood and have been reconstituted with purified proteins *in vitro* (Banjade & Rosen, 2014; Case et al., 2019). But how the WAVE complex organizes into lines is not understood and has not proven amenable to biochemical reconstitutions (Z. Chen et al., 2010; Koronakis et al., 2011; Lebensohn & Kirschner, 2009).

Here, we investigate the rules of the WAVE complex organization through *in vivo* biochemistry. With quantitative fluorescent imaging and analysis, we demonstrate that the WAVE complex forms linear arrays on cell membranes as expected for an NPF that builds sheet-like actin networks. To further investigate the biophysical properties of the WAVE complex linear arrays in a cellular context, we used single-molecule tracking in combination with fluorescent molecular counting and found that the WAVE complex forms solid-like linear arrays with a multilayered organization pattern. Moreover, the WAVE complex linear arrays are always associated with negative membrane curvature in the native cellular context. To test how membrane curvature affects WAVE complex assembly, we used nanobeads, nanopatterns, and cell compression to manipulate membrane curvature of the cells and found that negative membrane curvature is both sufficient and necessary for WAVE complex assembly on cell membranes. We propose a model of WAVE complex assembly as a multilayered linear array governed by negative membrane curvature that forms a template for lamellipodial formation.

## Results

### WAVE complex forms linear arrays on cell membranes

To build a sheet-like lamellipod, we expect the WAVE complex to organize into a linear array on cell membranes (**Figure 1**). In a migratory cell, the WAVE complex propagates as a “wave” pattern at the tips of lamellipodia with expanding and contracting zones of WAVE complex accumulation marking the expansion and contraction of the lamellipod (**Figure 2 B, C, and Video 1**). These “wave” patterns arise from an excitable feedback network composed of WAVE complex self-recruitment activated by Rho GTPases and delayed inhibition from actin polymerization that strips the WAVE complex off the plasma membrane (Millius et al., 2012; Pipathsouk et al., 2021; Weiner et al., 2007). The Rho GTPase Rac (assayed via the PBD [p21-binding domain] of PAK1 [p21-activated kinase 1]) broadly distributes at the front of the cell and spans a more permissive zone compared to the structured patterns of the WAVE complex, suggesting a self-organization propensity of the WAVE complex (**Supplementary Figure S1 A– D**; Pipathsouk et al., 2021; Weiner et al., 2007).

**Figure 2.**
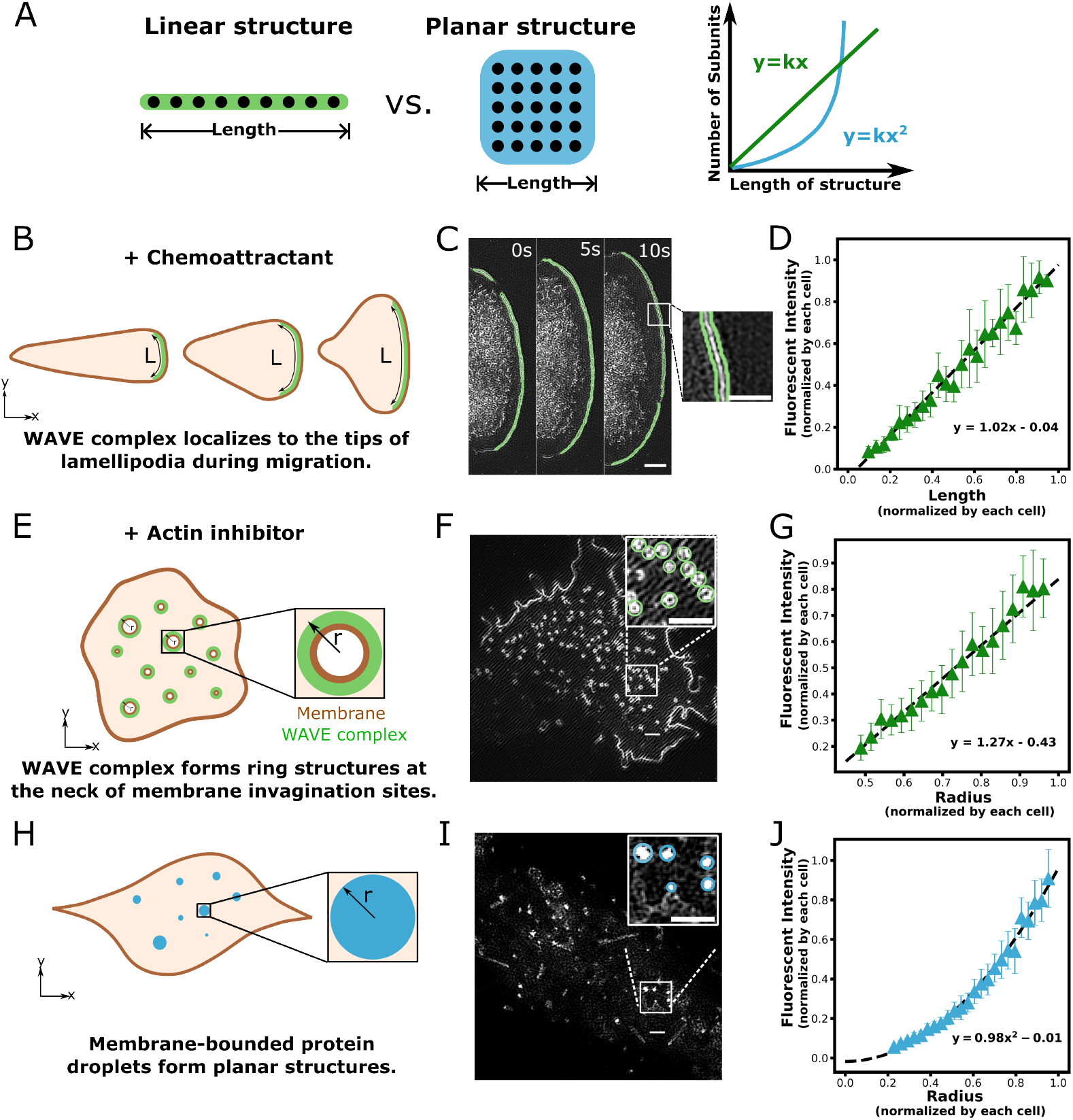
The WAVE complex is arrayed linearly on cell membranes. Here we sought to use quantitative fluorescence microscopy to probe the spatial organization of the WAVE complex in cells. (**A**) For a linear arrangement of subunits, we expect a linear relation between the length of the structure and the number of subunits (green). For a planar arrangement of subunits, we expect a quadratic relation between the length of the structure and the number of subunits (blue). (**B**) WAVE complex forms linear structures at the protruding tips of lamellipodia. (**C**) eGFP-Sra1 labeled WAVE complex in a migrating neutrophil-like HL-60 cell. Green contours indicate the region of the lamellipod used for WAVE complex quantification. Zoom in panel shows the inset within the magnified lamellipodia region. TIRF-SIM imaging; scale bar: 2 μm and 1 μm (inset). (**D**) WAVE complex enrichment at the tips of the lamellipodia scales linearly with lamellipodial length, suggesting an underlying linear organization of the WAVE complex. n = 331 lamellipodia from 32 cells acquired in three experiments. (**E**) WAVE complex forms nanoring structures in the absence of actin cytoskeleton. (**F**) eGFP-Sra1 labeled WAVE complex nanorings in an HL-60 cell treated with 0.5 μM actin inhibitor latrunculin B (latB). Upper right shows the white boxed inset with green circles indicating the quantitated WAVE complex nanorings. TIRF-SIM imaging; scale bars: 1 μm. (**G**) WAVE complex nanorings scale linearly with radius, suggesting an underlying linear organization of the WAVE complex. n = 5217 nanorings from 38 cells acquired in four experiments. (**H**) Membrane-bounded protein droplets (Fyn-HoTag3) adapted from the SPARK-ON system developed by Chung et al. (2023) form planar structures on cell membranes (a positive control for planar protein arrays). (**I**) Expression of Fyn-HoTag3 in HEK293T cells. Upper right inset shows magnified ROI with blue circles showing the quantified regions. TIRF-SIM imaging; scale bars: 1 μm. (**J**) The membrane-tagged SPARK-ON system exhibits a quadratic relation to radius, as expected for a planar structure. n = 407 puncta from 30 cells acquired in three experiments.

To quantify the native accumulation patterns of the WAVE complex on cell membranes, we used fluorescent quantification to measure the relation between the number of subunits and the pattern of the structure. A linear array exhibits a linear relation between the length of the array and the number of subunits, while a planar structure exhibits a quadratic relation between the length of the pattern and the number of subunits (**Figure 2 A**). To visualize the WAVE complex enrichment patterns on cell membranes, we expressed eGFP-Sra1 (one of the WAVE complex subunits) in neutrophil-like HL-60 cells and imaged them with total internal reflection fluorescence–structured-illumination microscopy (TIRF-SIM). Quantification of the fluorescent intensities of the WAVE complex vs. the lengths of the “wave” patterns at the tips of lamellipodia revealed a linear relation between them (**Figure 2 D**). To study the intrinsic WAVE complex accumulation patterns freed from the constraints of the actin cytoskeleton, we used latrunculin B (latB) to deplete actin polymers. Under these conditions, the WAVE complex forms nanoscale ring structures (**Figure 2 E and F, Video 1**; Pipathsouk et al., 2021). As for lamellipodia, the fluorescent intensity of the WAVE complex also scales linearly with the radius (length) of the intrinsic nanoring structures in cells treated with latB (**Figure 2 G**). This linear relation between the fluorescent intensity (that indicates the number of monomers) and the length of the pattern suggests an underlying linear organization of the WAVE complex. As a control for a planar structure (comparable to N-WASP focal droplets), we adapted the SPARK-ON system that drives protein separation in the cytosol through the multivalent interactions between HoTag3 and HoTag6 (Chung et al., 2023). By adding a membrane Fyn tag (Xia & Götz, 2014) to one of the SPARK-ON system components, HoTag3, we found that it can self-oligomerize when targeted to the cell membrane in HEK293T cells even in the absence of HO-Tag6 (**Figure 2 H and I, Supplementary Figure S1 E**), possibly due to the high local concentration of HoTag3 upon membrane recruitment. Unlike the WAVE complex that forms linear patterns on cell membranes, HoTag3 exhibits a quadratic relation between the fluorescent intensities and the radius of the pattern (**Figure 2 J**) and a linear relation to the area (**Supplementary Figure S1 F**), suggesting the formation of a planar structure.

### The WAVE complex is a core component of a linear array with a solid-like organization pattern

The WAVE-related NPF N-WASP undergoes liquid-liquid phase separation on cell membranes (Banjade & Rosen, 2014; Case et al., 2019). In contrast, the mechanism by which the WAVE complex forms linear arrays in cells is not known; purified WAVE complex does not assemble on its own (Z. Chen et al., 2010). We envisioned two possibilities for the linear organization of the WAVE complex –— either the WAVE complex binds to a pre-existing linear protein scaffold, like microtubule-associated proteins (MAP) binding to microtubules (Bodakuntla et al., 2019), or the WAVE complex could represent a core component of a linear array, like tubulin subunits in microtubules. To differentiate between these possibilities, we titrated the expression level of the WAVE complex in cells. If the WAVE complex is associated with an independently-formed protein scaffold, the WAVE complex concentration per unit length, which we measure by unit length fluorescent intensity on the scaffold, should scale within the WAVE complex expression level (**Figure 3 A**). In contrast, if the WAVE complex is a core component of a linear array, the WAVE complex should exhibit invariant concentration per unit length (**Figure 3 B**). We titrated WAVE complex abundance via Hem1-eGFP expression in Hem1 KO HL-60 cells. Disruption of any WAVE complex subunit destabilizes the remaining subunits of the WAVE complex (Innocenti et al., 2004; Mendoza, 2013), resulting in loss of lamellipodia formation in HL-60 cells (Graziano et al., 2019). Reintroduction of Hem1-eGFP in Hem1 KO cells titrates the WAVE complex from undetectable to endogenous levels of expression. We rescued the WAVE complex formation by expressing Hem1-eGFP in Hem1 KO cells followed by latB treatment to focus on the intrinsic WAVE complex organization without the complication of feedback inhibition from actin cytoskeleton (Mehidi et al., 2021; Pipathsouk et al., 2021; Weiner et al., 2007). We found that the WAVE nanoring structures in the absence of actin cytoskeleton exhibit invariant fluorescent intensity per unit length at all levels of Hem1-eGFP expression (**Figure 3 C and D, Supplementary Figure S2 A**), consistent with the idea that the WAVE complex is a core component of the linear array rather than associating with a pre-existing scaffold.

**Figure 3.**
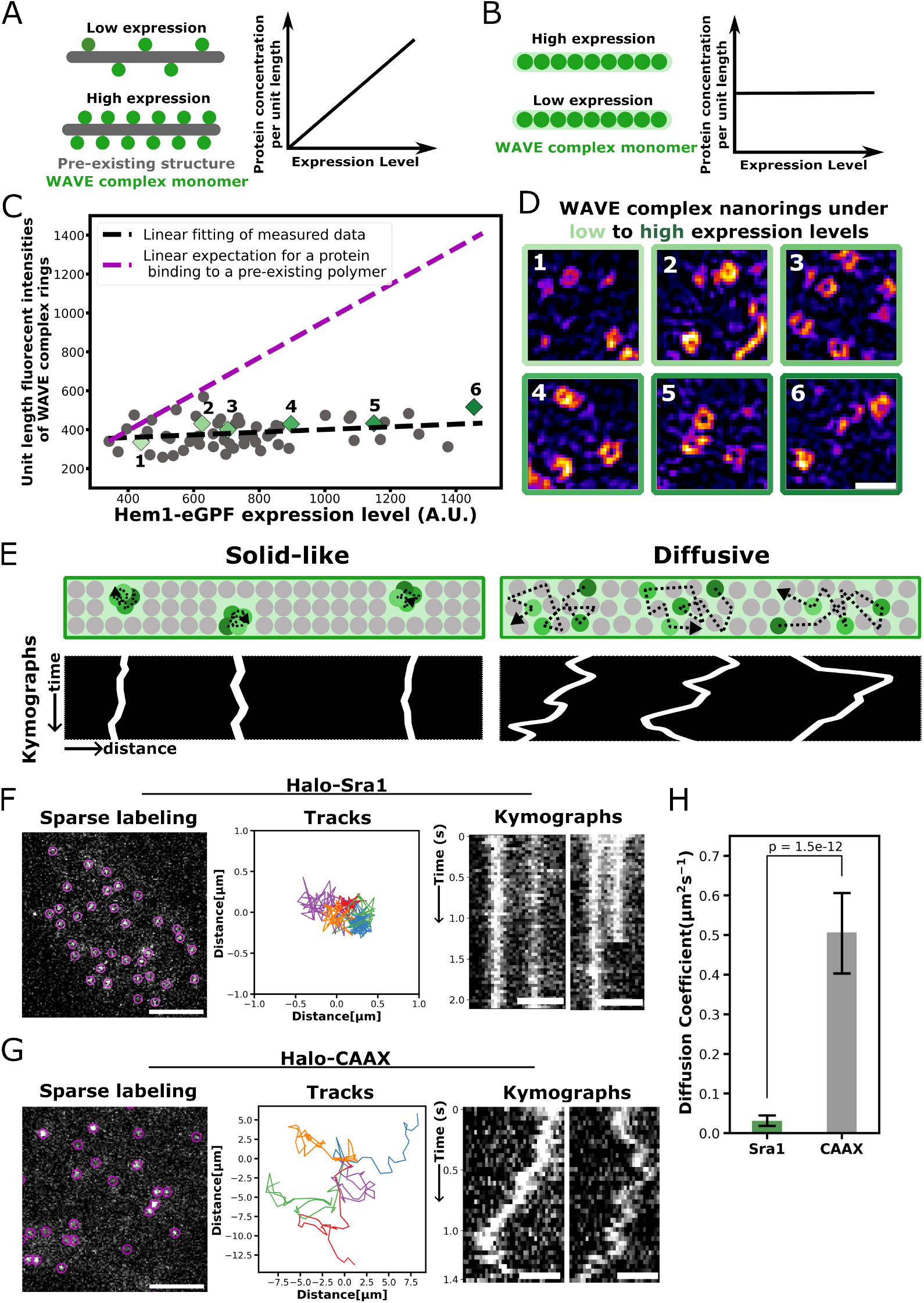
The WAVE complex is a core component of a linear array with a solid-like organization. (**A**) For proteins that bind to a pre-existing linear structure (like MAPs binding to microtubules), the protein concentration per unit length scales with the expression level of the protein. (**B**) Proteins that are core components of a linear structure should exhibit an invariant concentration per unit length regardless of expression level (like tubulin monomers forming microtubules). (**C**) Titrating the WAVE complex abundance via Hem1-eGFP expression in Hem1 KO cells shows that the WAVE complex exhibits an invariant concentration per unit length, suggesting that the WAVE complex is a core component of the linear array in which it is organized. Each dot corresponds to the average unit length intensity of WAVE complex nanorings (y axis; TIRF-SIM imaging) and corresponding Hem1 expression level (x axis; Epi-fluorescence imaging) in each cell, n=58 cells from three experiments. Examples of WAVE complex rings for the green diamond dots are shown in (**D**). Magenta line shows the linear relation between the unit length intensity of the WAVE complex rings and Hem1-eGFP expression level that would be expected for a protein that binds a pre-existing array. (**D**) Example images of the WAVE complex rings corresponding to each diamond dot in C with different shades of greens indicating different expression levels. TIRF-SIM imaging; scale bars: 0.5 μm. (**E**) We employed single-molecule tracking to differentiate between two different potential modes of protein organization structures—solid-like arrays (like actin filaments or microtubules) and diffusive protein associations (like liquid-liquid protein condensates such as N-WASP). Bottom shows the mock kymographs of the single particle dynamics in each mode. (**F**) Single-molecule tracking data of the Halo-tagged Sra1 cell line shows that the WAVE complex remains stationary in the latB-treated cells. Left: sparse labeling of the Halo-Sra1 HL-60 cells. Each magenta circle annotates a single Sra1 subunit of the WAVE complex. TIRF imaging; scale bar: 5 μm. Middle: selected tracks of Sra1 subunits. Right: kymographs of the Sra1 subunit indicating a solid-like organization of WAVE complex within the linear array. Scale bar: 2 μm. (**G**) Single-molecule tracking data of the Halo-tagged CAAX cell line show the dynamic and diffusive behavior of the membrane-binding CAAX protein in latB-treated HL-60 cells. TIRF imaging; scale bar: 5 μm. Left: sparse labeling of the Halo-CAAX HL-60 cells. Each magenta circle annotates a single CAAX protein. Middle: selected tracks of single CAAX proteins. Right: kymographs showing the diffusive nature of CAAX in the cell membrane. Scale bar: 2 μm. (**H**) Diffusion coefficient of the WAVE complex (green; n = 1763 from 12 cells) vs. the CAAX protein (grey; n = 2157 from 24 cells). P = 1.5e-12 by an unpaired two-tailed t-test.

Next, we investigated the modes of WAVE complex subunit organization on cell membranes. Those linear arrays could either be solid-like (like actin filaments or microtubules) or diffusive (like N-WASP). If the WAVE complex forms solid-like structures, each subunit within the linear array should remain relatively stationary to other subunits rather than exhibiting a diffusive behavior (**Figure 3 E**). To differentiate between these two modes, we employed single-molecule tracking to study the subunit dynamics within the WAVE complex linear arrays on cell membrane. We expressed Halo-tagged Sra1 in HL-60 cells and labeled the cells with two HaloTag ligand dyes, a high concentration of JF646 for full labeling of the WAVE complex (**Supplementary Figure S2 B and C**) and a lower concentration of JF549 (sparse labeling) for single-molecule tracking (**Figure 3 F**). As a control for a protein that is diffusive in the membrane, we expressed Halo-tagged membrane binding motif CAAX in HL-60 cells. Cells were treated with actin inhibitor to avert cytoskeletal-applied forces and actin-based recycling (Mehidi et al., 2021; Weiner et al., 2007) and then imaged with TIRF microscopy at a rate of ~26 frames per second (**Supplementary Figure S2 D**). The single molecule traces indicate that the WAVE complex monomers in the linear WAVE complex arrays are mostly stationary and minimally diffusing, whereas the CAAX-tagged proteins exhibit significant lateral diffusion (**Figure 3 F–H, S2 B and C, Video 2**). These data suggest that the WAVE complex linear arrays adopt a solid-like organization pattern, similar to protein polymers like actin filaments or microtubules.

### The WAVE complex linear array exhibits a multilayered organization

The WAVE complex solid-like linear array could either adopt a single-layered arrangement similar to actin filaments (Katsuno et al., 2015) or a multi-layered arrangement similar to microtubules (Roostalu & Surrey, 2017). To investigate the subunit arrangement of the WAVE complex linear array, we performed molecular counting of the WAVE complex linear arrays on cell membranes. If the WAVE complex is forming a single-layered array, it would necessitate at least 50-125 copies of the WAVE complexper micron given the dimension (200Å × 110Å × 80Å) of a single WAVE complex (**Figure 4 B**; Z. Chen et al., 2010). To determine the actual number of the WAVE complexes within the linear array, we leveraged fluorescent standard candles to calculate the number of fluorescent molecules using the microscopically-measured fluorescent intensities to construct a fluorescence-based standard curve (Akamatsu et al., 2020; Hsia et al., 2016). The membrane-tagged fluorescent candles with 12, 60, or 120 copies of eGFP proteins were expressed in HL-60 cells (**Figure 4 A**). We used TIRF-SIM to measure the fluorescence intensities of the standard candles in cells that were chemically fixed to limit movement of the candles on cell membrane (**Figure 4 A**). With this data, we constructed the fluorescence-based standard curve that confirms good linearity between the fluorescent intensities and the number of fluorescent molecules in each standard candle (**Figure 4 A**).

**Figure 4.**
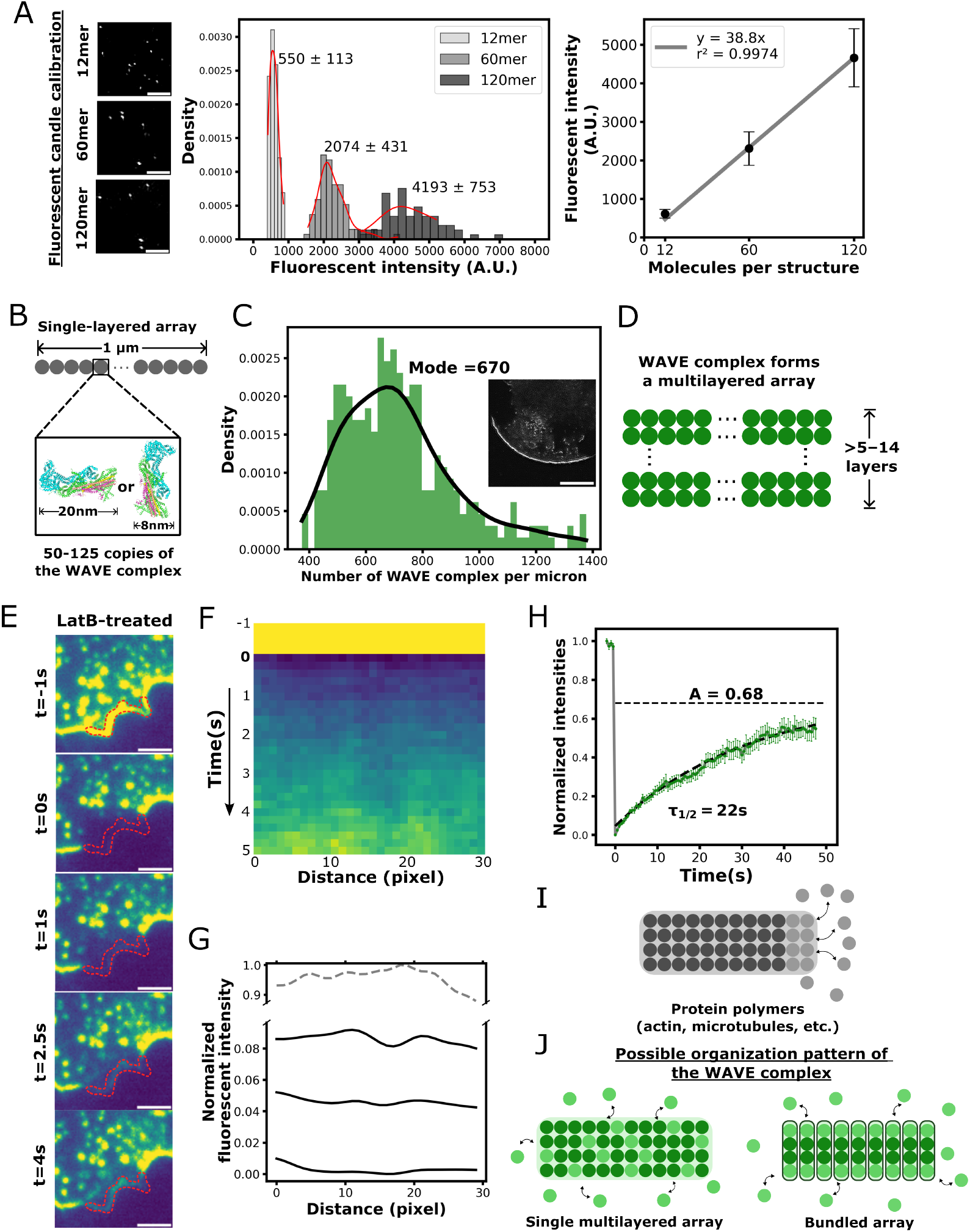
The WAVE complex linear array has a multilayered organization. To investigate the basis of WAVE complex subunit organization in linear arrays in cells, we performed molecular counting of the number of WAVE complexes in lamellipodia using the fluorescent standard candles developed by Hsia et al., 2016 and Akamatsu et al., 2020 that contain known numbers of eGFPs. Left: Expression of the fluorescent standard candles in HL-60 cells. Cells were fixed with glutaraldehyde. TIRF-SIM imaging; scale bar: 1 μm. Middle: Histograms of the fluorescent distributions of 12-mer (n = 58 from 11 cells), 60-mer (n = 85 from 9 cells), and 120-mer (n = 64 from 10 cells) candles. Red lines show the kernel density estimation of each distribution. Right: Fluorescence-based standard curve showing the relation between the fluorescent intensities and the numbers of fluorescent molecules. Line shows linear fit through zero. **(B)** If the WAVE complex is adopting a single-layered arrangement like actin filaments, this would require 50-125 copies of the WAVE complex to make an array of 1 μm, based on PDB 3p8c by Z. Chen et al., 2010. (**C**) Histogram of the number of WAVE complexes with a mode of 670 per micron in lamellipodia (n = 292 from 27 cells over three experiments). This number was calculated using the standard curve in panel B, corrected by the fixed v.s. unfixed fluorescence ratio (**Supplementary Figure S3 A**) and the percentage of eGFP-tagged WAVE complex as determined by Western Blot (**Supplementary Figure S3 B**). Black line shows the kernel density estimation of the distribution. Middle right inset shows the TIRF-SIM live imaging of the eGFP-Sra1 labeled WAVE complex at the tips of lamellipodia (scale bar: 5 μm) in living cells using the same imaging setup as the standard candles. (**D**) Based on these data, the WAVE complex linear array must adopt a multilayered arrangement at least 5-14 layers thick (depending on the orientation of the WAVE complex). (**E**) We employed FRAP of the WAVE complex array to study its turnover dynamics. Bleached WAVE complex begins recovering within a couple of seconds in latB-treated HL-60 cells. Ring-TIRF imaging; scale bar: 2 μm. (**F**) Kymographs showing the recovery of the WAVE complex fluorescent signal within 5 s after photobleaching in latB-treated HL-60 cells. Data represent averages from 12 bleached sites from 12 cells. (**G**) Line scans of the kymographs at different time points show uniform recovery within the WAVE complex array. (**H**) Quantification of the WAVE complex fluorescence recovery after photobleaching in latB-treated cells. The WAVE complex shows partial recovery with a ratio of 68%. N = 12 cells. (**I**) Protein polymers (actin or microtubules) typically exchange at the end of the array (and not in the interior of the array). (**J**) Based on the FRAP data, we envision two possible organization patterns of the WAVE complex. Left: a single multilayered array that permits internal subunit exchanges. Right: a set of bundles where subunit exchanges happen at the end of each bundle.

Next, we used the eGFP-Sra1 subunit of the WAVE complex to calculate the absolute abundance of WAVE complex at the cell membrane in HL-60 cells. The total expression level of the WAVE complex in each cell was not affected by eGFP-Sra1, as any subunit not incorporated into the WAVE complex is degraded (Graziano et al., 2019; Pipathsouk et al., 2021; Weiner et al., 2006). We quantified the fluorescent intensities of the eGFP-Sra1 at the tips of lamellipodia. Correcting for fluorescence loss after fixation and the tagged ratio of the Sra1 subunits (**Supplementary Figure S3 A and B**) yielded a mode of 670 copies of the WAVE complex per micron at the tips of lamellipodia (**Figure 4 C**). This value of 670 is much more than the copies of WAVE complex required to make a single-layered array (**Figure 4 B**), suggesting that WAVE complex form a multilayered array at least 5-14 layers thick at the tips of lamellipodia (**Figure 4 D**). Similarly, we also performed molecular counting of the WAVE complex nanorings in cells treated with latB and determined a mode of 260 copies of the WAVE complex within the nanoring structures, suggesting that the WAVE complex also adopts a multilayered organization pattern in the nanorings (**Supplementary Figure S3 C and D**).

To investigate the mechanism of WAVE complex linear array assembly, we used fluorescence recovery after photobleaching (FRAP) to study the turnover dynamics of the WAVE complex on cell membranes. The WAVE complex at the tips of lamellipodia in migrating cells exhibit rapid turnover dynamics with a half-life of 6.3 s and complete exchange along the length of the array (**Supplementary Figure S3 E and Video 3**; Mehidi et al., 2021; Weiner et al., 2007). When treated with actin inhibitor, the turnover dynamics of the WAVE complex are decreased to 22 s with about one-third of the WAVE complex molecules maintaining their position within the linear array (**Figure 4 H**). The rest exchanging molecules exhibit a uniform recovery pattern along the bleached region (**Figure 4 E–G and Video 4**), indicating that the molecule exchange is happening between the array and the cytosolic pool with minimal diffusion within the array itself (**Figure 3 F–H**). This turnover behavior is notably distinct from actin or microtubule polymers where subunits only exchange at the end of the protein array (**Figure 4 I**; Katsuno et al., 2015; Smith et al., 2013; Vorobjev et al., 1999). This uniform but partial recovery pattern suggests that the WAVE complex could either be 1) a single multilayered linear array like a microtubule but permits molecule exchange within the array as microtubules do under certain conditions (Kuo et al., 2022), or 2) a bundled array like actin in stress fibers (Tojkander et al., 2012) where molecular exchanges occur at the end of each short polymer within the array (**Figure 4 J**).

### Negative membrane curvature is both sufficient and necessary for WAVE complex membrane association

Some multilayered polymers like microtubules can polymerize into a curved parallel array where their lateral extension of polymerization is constrained by the intrinsic curvature of the polymer (Chrétien et al., 1995; Nogales & Wang, 2006). Is curvature playing a similar role for the assembly of the multilayered WAVE complex linear arrays? Consistent with this idea, there is a characteristic membrane curvature seen for WAVE complex recruitment in cells. WAVE complex localizes to the tips of the lamellipodia with negative curvature of a radius of 65 nm in the axis of protrusion (x-z plane) and little-to-no curvature along the lamellipod (x-y plane) (**Figure 5 A**; Pipathsouk et al., 2021; Schmeiser and Winkler, 2015). For cells treated with actin inhibitor, WAVE complex forms nanorings localized to the neck of membrane invagination sites that exhibit saddle curvature, consisting of a protruding negative curvature of a radius of 65 nm perpendicular to the invagination neck (x-z plane) and a radius curvature of 115 nm around the invagination neck (x-y plane) (**Figure 5 B**; Pipathsouk et al., 2021). Despite dramatic differences in the overall membrane morphology, the 65 nm radius of negative curvature is preserved between both of these contexts. These data suggest that this negative membrane curvature could be a key permissive factor in WAVE complex association on cell membranes.

**Figure 5.**
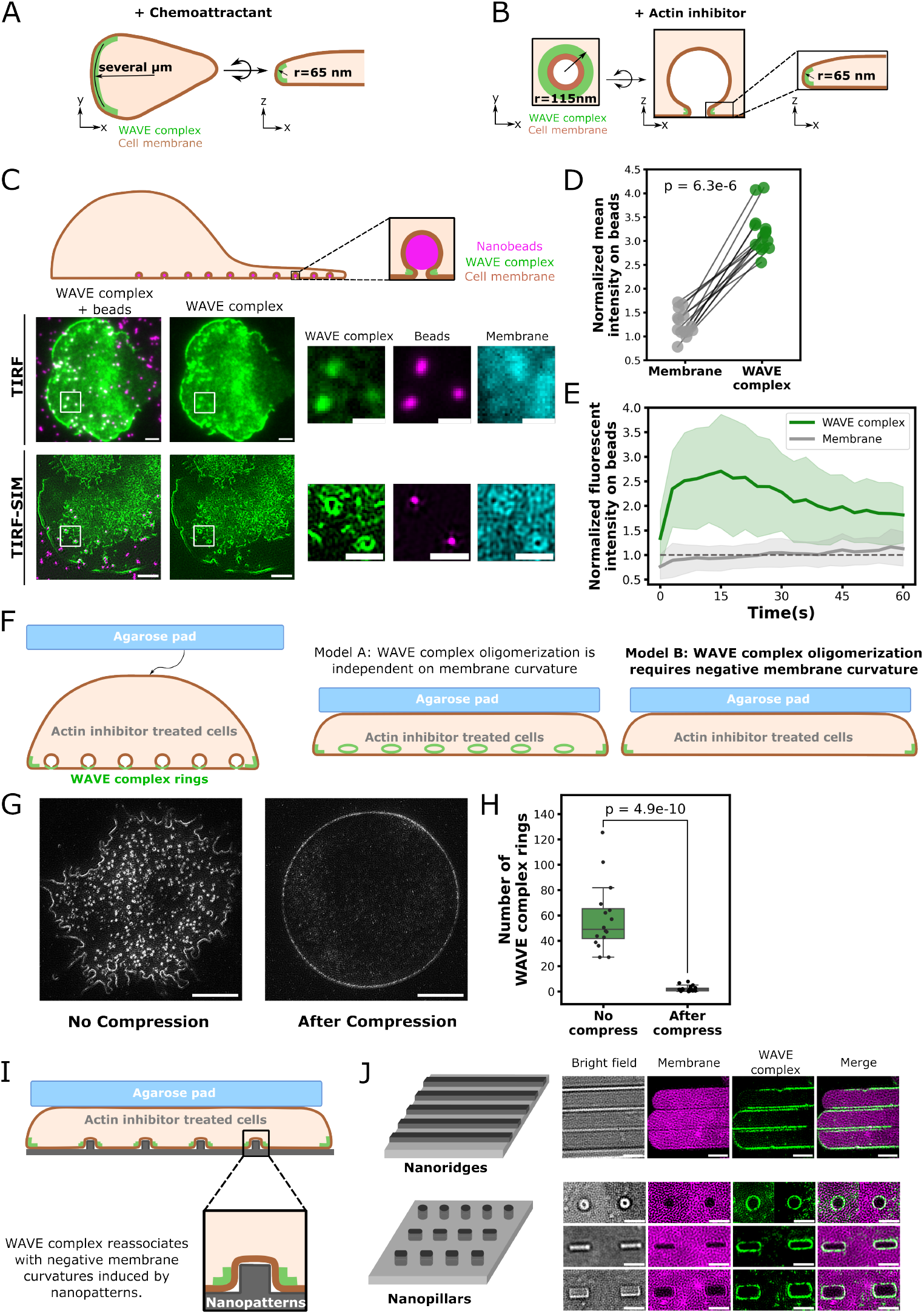
Membrane curvature is both sufficient and necessary for WAVE complex membrane association. (**A**) The WAVE complex localizes to the tips of lamellipodia that exhibit a radius of negative curvature (x-z plane) of 65 nm and little to no curvature along the lamellipod (x-y plane). (**B**) The WAVE complex nanorings in cells treated with actin inhibitor localize to the neck of the membrane invagination site with a radius of negative curvature (x-z plane) of 65 nm and positive curvature (x-y plane) of 115 nm. (**C**) To test if membrane curvature is sufficient for WAVE complex membrane association, we used nanobeads to induce negative membrane curvatures in migrating HL-60 cells (top). WAVE complex (eGFP-Sra1; green) colocalizes with and forms ring structures around the 200 nm beads (magenta). Insets shown on right. Scale bars: 2 μm and 1 μm (insets). (**D**) Quantification of the WAVE complex intensity on beads compared to membrane marker (Cell Mask DeepRed). WAVE complex shows significantly increased signals on beads, showing the ability of negative membrane curvature to induce WAVE complex membrane recruitment. P = 6.3e-6 by a paired two-tailed t-test on the mean normalized WAVE complex/membrane intensity at all beads in each cell. n = 14 cells. (**E**)The WAVE complex (green) shows persistent association with beads over 60 s. n = 120 beads from 14 cells. (**F**) To test if membrane curvature is necessary for WAVE complex membrane association, we compressed the latB-treated HL-60 cells with an agarose pad to iron out membrane invagination sites. Persistence of WAVE complex nanorings would suggest that membrane curvature is dispensable for WAVE complex membrane association in linear arrays (model A). Disappearance of WAVE complex nanorings following compression would suggest that membrane curvature is necessary for WAVE complex membrane association in linear arrays (model B). (**G**) eGFP-Sra1 HL-60 cells before(left) and after (right) compression. The vast majority of the WAVE complex nanorings disappear following compression. The WAVE complex arrays are only maintained at the rim of the cell where the negative membrane curvature persists following compression. TIRF-SIM imaging; scale bars = 5 μm. (**H**) WAVE complex nanoring abundance before (n = 16 cells) and after compression (n = 18 cells). P = 4.9e-10 by an unpaired two-tailed t-test. (**I**) Rescuing WAVE complex membrane association after compression with nanopatterns that can induce negative membrane curvature. (**J**) WAVE complex (green) reassociates on cell membrane(magenta) with negative curvatures induced by nanopatterns even in the presence of the agarose-based compression. TIRF-SIM imaging; scale bar = 2 μm.

To test the role of negative membrane curvature in WAVE complex linear assembly, we first investigated whether negative curvature is sufficient for WAVE complex membrane association. We exposed migrating cells to polystyrene beads of 200 nm that could induce membrane invaginations with negative curvature (**Figure 5 C**). As eGFP-Sra1 tagged HL-60 cells migrate over these beads, the WAVE complex forms ring structures at the neck of membrane invagination sites (**Figure 5 C, Video 5 and 6**). The WAVE complex signal on beads is persistent on beads over time and significantly higher than the membrane dye signal (**Figure 5 D and E, S4 A and B**), indicating that the increase in WAVE signal is not due to excess membrane accumulation but rather reflects WAVE complex accumulation at the negative membrane curvature induced by the beads. These data suggest that negative curvature is sufficient to induce WAVE complex membrane association in the context of a migrating cell.

Next, we tested whether negative membrane curvature is necessary for WAVE complex membrane association by inhibiting negative membrane curvature. The native negative curvature that proved most amenable to manipulation was the membrane invaginations at the ventral surface of latB-treated cells. We overlaid cells with a precast agarose pad to induce compression and iron out the membrane invagination sites. If negative curvature is required for WAVE complex membrane association, disturbing membrane invagination sites should interfere with the WAVE recruitment to nanorings (**Figure 5 F**). Indeed, the WAVE complex nanorings disappear following cell compression (**Figure 5 G and H**). Because the WAVE complex persists at the rim of the cell where the negative curvature is maintained (**Figure 5 G**), it is unlikely that compression exerts its effects by blocking the pstream regulators WAVE complex recruitment.

Finally, we tested whether WAVE complex membrane association can be rescued during compression by reintroducing negative membrane curvature to cells. For this purpose, we provided latB-treated cells with nanopatterns at the ventral surface that could sustain negative membrane deformations following compression with the agarose pad (**Figure 5 I**; Cail et al., 2022). Under these conditions, WAVE complex forms linear arrays around the regions of micropattern-induced negative curvature (**Figure 5 J, S4 C**). Moreover, WAVE complex can also be recruited to regions of negative curvature around nanopatterns in migratory cells (**S4 D, Video 7**). Together, these data suggest that the negative membrane curvature plays a key role in controlling WAVE complex association on cell membranes.

## Discussion

The WAVE complex is an actin regulator that organizes many different cellular behaviors, including chemotaxis, macropinocytosis, phagocytosis, and cell integrity (Davidson et al., 2018; King & Kay, 2019; Maddugoda et al., 2011; Pipathsouk et al., 2021; Seastone et al., 2001). These processes all involve flat, sheet-like actin-based protrusions (Humphreys et al., 2016; Mylvaganam et al., 2021; Veltman et al., 2016a; Yang et al., 2021), but how the WAVE complex instructs this pattern of protrusion is not well understood. Since the WAVE complex does not exhibit these behaviors in cell-free biochemical reconstitutions (Z. Chen et al., 2010; Koronakis et al., 2011; Lebensohn & Kirschner, 2009), our work leverages *in vivo* biochemistry approaches to understand the logic of lamellipodia formation. We reveal a linear self-organization pattern of the WAVE complex at the cell membrane that could form a template for these flat, sheet-like actin networks (**Figure 1 and Figure 2**). Unlike N-WASP that undergoes phase separation to form focal liquid-like protein condensates (Banjade & Rosen, 2014; Case et al., 2019), WAVE complex monomers form ordered linear arrays in which the subunits exhibit a fixed position relative to one another (**Figure 3**), suggesting linear polymer-like structures at the cell membrane (**Figure 6 A**). Similar to microtubules, these linear arrays of the WAVE complex exhibit a multilayered organization (**Figure 4**) whose lateral extent (and thus the width of the polymer) appears to depend on the curvature of the underlying polymer (**Figure 6 A**; Chrétien et al., 1995; Nogales and Wang, 2006). In the case of the WAVE complex, membrane curvature is necessary and sufficient for WAVE complex membrane recruitment (**Figure 5**).

**Figure 6.**
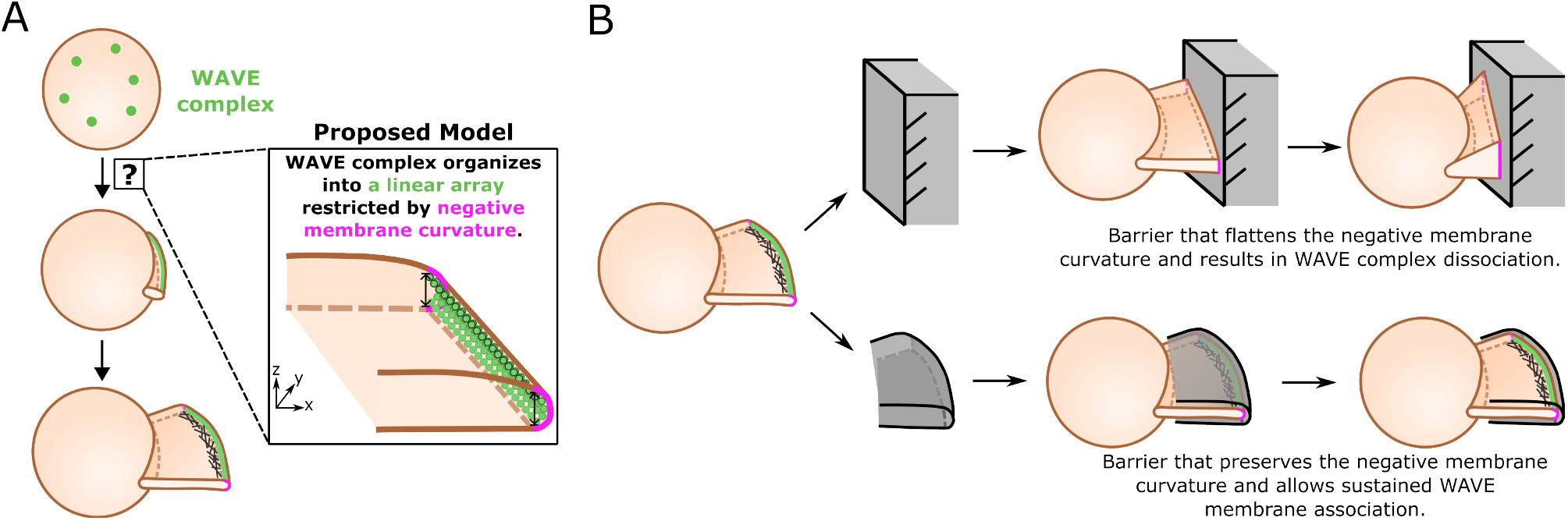
The WAVE complex organizes into a multilayered linear array at negative membrane curvature. (**A**) Our proposed model of how the WAVE complex organizes into linear arrays in cells: a multilayered solid-like linear array whose lateral extension is restricted by negative membrane curvature. (**B**) Our model highlights the potential role of membrane curvature in barrier avoidance. When a cell runs into a barrier (upper), the negative curvature at the tips of the lamellipod is disturbed, resulting in WAVE complex dissociation and cell stalling. For barriers that preserve the shape of the lamellipod (lower), cells fail to recognize the barrier and have sustained WAVE complex association at the negative membrane curvature provided by the barrier (**Figure 5 E**).

The WAVE complex plays a central role in lamellipodia formation. For neutrophils and many other cells, depletion of the WAVE complex abolishes cells’ ability to build lamellipodia (Graziano et al., 2019; Law et al., 2013; Leithner et al., 2016; Whitelaw et al., 2020). The WAVE complex stimulates activation of the Arp2/3 complex, but the Arp2/3 complex is not required for WAVE’s ability to generate lamellipodia (Buracco et al., 2022; Pipathsouk et al., 2021). Neither the Arp2/3 complex or actin assembly are needed for the WAVE complex to form linear arrays at the cell membrane (Pipathsouk et al., 2021). Our work suggests the WAVE complex organization into linear arrays could be a key ingredient for its role in lamellipodial formation. Here the WAVE complex also plays a central role — the WAVE complex is a core component of these linear arrays rather than decorating a pre-existing polymer (**Figure 3 A–D**). However, the WAVE complex isn’t self-organizing on its own. In the presence of upstream activators, motility mix, and complete cytosol, the WAVE complex initiates actin polymerization but fails to form any high-order oligomers (Koronakis et al., 2011; Lebensohn & Kirschner, 2009), nor does the WAVE complex oligomerize on its own (Z. Chen et al., 2010). The additional inputs that enable WAVE complex assembly into linear arrays are not known. One possibility is that these previous assays fail to provide the strong negative membrane curvature that we have found to be required for WAVE complex membrane recruitment (**Figure 5**); previous assays have used positively curved surfaces like vesicles or lipid-coated beads (Humphreys et al., 2016; Lebensohn & Kirschner, 2009) or planar lipid bilayers (Banjade & Rosen, 2014; Lebensohn & Kirschner, 2009). In addition, the higher order organization of the WAVE complex could require additional membrane-bound partners, including those containing WRC interacting receptor sequence binding proteins (B. Chen et al., 2014) that have been lacking in existing reconstitutions.

In addition to the membrane-bound upstream activators of the WAVE complex such as Rac, Arf, and phosphoinisitides (Koronakis et al., 2011; Oikawa et al., 2004; Suetsugu et al., 2006), our work suggests an additional requirement for cell geometry in WAVE complex patterning, in particular strong negative membrane curvature. This curvature dependence could help explain some of the emergent behaviors of cell migration. For example, when cells undergo a collision with an obstacle, they cease migration and change directions (Roycroft & Mayor, 2016; Weiner et al., 2007; Yamada & Sixt, 2019). This barrier avoidance could be achieved through several possible mechanisms, including forces acting at the leading edge (Diz-Muñoz et al., 2016; Nakamura, 2024; Prass et al., 2006; Saha et al., 2018; Uray & Uray, 2021) or the nucleus (Lomakin et al., 2020; Long & Lammerding, 2021; Venturini et al., 2020), cell stalling (Brunner et al., 2006) and/or perturbation of cell shapes and curvatures (**Figure 6 B**; Sitarska et al., 2023). If membrane curvature is a primary input to barrier avoidance, then the barriers won’t be recognized if they don’t change the morphology (in particular the strong negative curvature) of the lamellipodia. Consistent with this idea, lamellipodia do not stall at the necks of membrane invaginations or nanopatterns, potentially because the overall negative curvature of the lamellipodia is preserved at these barriers(**Figure 5 F, Supplementary Figure S3 A and B, Video 5–7**). Membrane curvature could also enable cell repolarization after collision, as computational modeling shows that negative membrane curvature-sensing proteins that recruit the actin cytoskeleton suffice to guide cell reorientation and spreading by selectively aligning with the substrate (Sadhu et al., 2023).

How the WAVE complex recognizes negative curvature is not known. The WAVE complex does not have any well-characterized curvature-sensing motifs. It interacts with iRSp53, a negative-curvature sensing I-BAR protein that is dependent on its interactions with the WAVE complex to localize to the tips of lamellipodia. However, depletion of iRSp53 does not affect WAVE complex’s ability to recognize negative membrane curvature (Pipathsouk et al., 2021), and depletion of all four of the I-BAR proteins does not prevent lamellipodia formation (Pokrant et al., 2023). It is possible that the WAVE complex may interact with additional curvature-sensitive proteins (Frost et al., 2009; Hanawa-Suetsugu et al., 2019; S. Liu et al., 2015; Pykäläinen et al., 2011). Alternatively, assembly of WAVE complex into a linear array could present the monomers in a geometry that is amenable to curvature sensation. Other curvature sensing proteins, like septins or the BAR domain family, have been shown to exhibit dramatically different curvature preferences following oligomerization compared to the preferences of the isolated monomers (Bridges et al., 2016; Nepal et al., 2021).

Lamellipodia are common protrusive strauctures for a wealth of migratory and morphogenetic contexts (Fritz-Laylin et al., 2017; Ibarra et al., 2006; Kunda et al., 2003; Leithner et al., 2016; Pollard & Borisy, 2003; Rakeman & Anderson, 2006; Weiner et al., 2006; Yolland et al., 2019), suggesting a potential value of lamellipodia over other membrane structures. The broad actin networks that compose lamellipodia can generate greater protrusive forces compared to filopodia and ensure uniform advancing of the cell membrane (Dimchev et al., 2017; Gardel et al., 2010; Pipathsouk et al., 2021), while their flat and thin geometry could enable efficient barrier avoidance, spreading along surfaces (Sadhu et al., 2023), and regulation of membrane reservoirs (Gauthier et al., 2011; Mueller et al., 2017). The flat, sheet-like membrane structures built by the WAVE complex are also involved in other morphogenetic processes. During macropinocytosis/phagocytosis, the WAVE complex assembles at the rim of the macropinocytic/phagocytic cups (King & Kay, 2019; Seastone et al., 2001; Veltman et al., 2016b). This discrete distribution depends on upstream biochemical activators like Ras and PIP3 (Buckley et al., 2020; Yang et al., 2021); our results could indicate that this focused accumulation could also be informed by the morphology of the macropinosome.

Our work further underscores the importance of NPF self-organization in instructing cell protrusion morphology. The default organization pattern for NPF-based actin assembly at membranes is filopodium-like protrusions (A. P. Liu et al., 2008). To generate different organizations of actin networks, *in vitro* biochemical reconstitutions have used the same NPF artificially arrayed into different patterns like points (which generate filopodial-like actin networks) or lines (which generate lamellipodial-like actin networks) (Boujemaa-Paterski et al., 2017; Carlier et al., 2003). In cells, different NPFs are used for different morphological structures, likely because these NPFs have different patterns of self-organization: N-WASP undergoes phase separation and forms focal protein droplet structures that orchestrate the fingerlike filopodia formation (Banjade & Rosen, 2014; Case et al., 2019), while our work reveals the key features of the WAVE complex linear array that instructs lamellipodial formation (**Figure 6 A**). In future work, it will be important to define how other patterns of actin regulator self-organization underly other actin structures including the cell cortex and stress fibers (Bovellan et al., 2014; Hotulainen & Lappalainen, 2006; Lehtimäki et al., 2021; Svitkina, 2020; Tojkander et al., 2012).

## Methods

### Cell culture

HL-60 cells were cultured in RPMI 1640 media supplemented with l-glutamine and 25 mM HEPES (MT10041CV; Corning) containing 10% (vol/vol) heat-inactivated fetal bovine serum (16140071; Gibco BRL). Cultures were maintained at a density of 0.2 to 1.0 million cells/mL at 37°C/5% CO2. HL-60 cells were differentiated with 1.3% (vol/vol) DMSO (358801; Santa Cruz Biotechnology) in culture media for 5 days before experiments. HEK293T cells were grown in DMEM (SH30243.01; Cytiva) containing 10% (vol/vol) heat-inactivated fetal bovine serum and maintained at 37°C/5% CO2.

### Transduction of HL-60 cells

HEK293T cells were seeded into 6-well plates (2.5 mL per well) and grown until about 70% confluent. For each well, 1.5 μg pHR vector (containing the appropriate transgene), 0.167 μg vesicular stomatitis virus-G vector, and 1.2 μg cytomegalovirus 8.91 vector were mixed and prepared for transfection using TransIT-293 transfection reagent (Mirus Bio) per the manufacturer’s instructions. After transfection, cells were grown for *~*48 hours, after which virus-containing supernatants were harvested and concentrated 30–40 fold using a Lenti-X Concentrator (Takara Bio) according to the manufacturer’s instructions. Lentivirus was used immediately or stored at −80 °C. HL-60 cells were transduced by overnight incubation of 0.32 million cells with 4 μg/mL polybrene and 125 μL of concentrated virus. Cells expressing desired transgenes and expression levels were isolated using FACS (FACS Aria2 or FACS Aria3; BD Biosciences). For the cells expressing the fluorescent standard candles, the lowest expression cells above the background were selected to avoid multiple candle aggregation (more than one candle in each spot).

### Transient transfection of HEK293T cells

HEK293T cells were seeded into 24 well glass bottom plates (P24-1.5H-N; Cellvis) and grown until about 70% confluent. For each well, 0.5 μg plasmid containing the appropriate transgene were used for transfection using TransIT-293 transfection reagent (Mirus Bio) according to the manufacturer’s instructions. After transfection, cells were grown for 24 hours before imaging.

### Plasmids

Plasmids were constructed using standard molecular biology protocols. DNA segments were PCR amplified and cloned into a pHR lentiviral backbone and driven by promoter from spleen focus-forming virus (SFFV) via standard Gibson assembly. The membrane-tagged Hotag3 construct was modified from Chung et al., 2023 by inserting a membrane-bounded Fyn tag into the N-terminus of the original coding sequence. The fluorescent standard candle constructs were modified from Akamatsu et al., 2020, by switching the fluorescent protein from tag2GFP to eGFP followed by subcloning into the pHR lentiviral backbone.

### Immunoblotting

Protein content from one million HL-60 cells was extracted by chilled TCA precipitation and resuspended in 2x Laemmli sample buffer. Protein samples were separated via SDS–PAGE, followed by transfer onto PVDF membranes (#1620177; BIO-RAD). Membranes were blocked at RT for 1 h in a 1:1 solution of TBS (20 mM Tris, 500 mM NaCl, pH 7.4) and Odyssey Blocking Buffer (#927-40000; LI-COR) followed by overnight incubation at 4 °C with primary antibodies in a solution of 1:1 TBST (TBS + 0.2% wt/vol Tween 20) and Odyssey Blocking Buffer. Membranes were then washed three times with TBST and incubated for 45 min at RT with secondary antibodies diluted 1:10,000 in 1:1 solution of Odyssey Blocking Buffer and TBST. Membranes were then washed three times with TBST, one time with TBS, and imaged using an Odyssey Fc (LI-COR). Primary antibodies Sra1 (rabbit; 1:200; #NBP2-16060; Novus), and secondary antibodies IRDye 680RD goat anti-mouse (926-68070; LI-COR) and HRP-Conjugated goat anti-rabbit (65-6120; Invitrogen) were used for this study.

### Cell preparation for imaging

#### Membrane labeling

Membrane labeling solution was made with CellMask Deep Red (Invitrogen) freshly diluted 1:500 in imaging media (Leibovitz’s L-15 [Gibco] with 2% FBS). 1 mL of differentiated HL-60 cells were spun down at 200x g for 3 min and resuspended in 500 μL of the labeling solution. Immediately after resuspension, cells were washed twice with imaging media before plating.

#### Sparse labeling cells for single molecule tracking

To label the Halo-Sra1 cells, 0.05 nM JF549 (GA1110; Promega) dye was mixed with 10 nM JF646 dye (GA1120; Promega). To label the Halo-CAAX cells, 0.001 nM JF549 with 10 nM JF646 were used. Cells were incubated with the dyes diluted in imaging media in a 37 °C/5% CO2 incubator for 20 min followed by two washes with imaging media.

#### Cell migration on fibronectin-coated coverslips

The desired number of wells in a 96-well plate (MGB096-1-2-LG-L; Matrical, Inc.) or an 8-well Lab-Tek II chamber (155409; Thermo Fisher Scientific) were coated with 100 μL or 200 μL of 40 μg/μL porcine fibronectin (prepared from whole blood) diluted in imaging media for 20 min at room temperature, followed by two to three times with imaging media. Differentiated cells (~0.8 million/μL) were resuspended in the imaging media of the same volume. 100 μL cells per well were used for a 96-well plate and 200 μL cells per well were used for an 8-well chamber. For cell migration on beads, 200 μL cells were mixed with 0.8 μL 0.2 μm red fluorescent (580/605) carboxylate-modified microspheres (F8887; Invitrogen) before plating. The imaging plate was then transferred to a 37°C/5% CO2 incubator for 15 min to allow cells to adhere. The plate was next transferred to the microscope, which had been preheated to 37°C for imaging. For chemoattractant stimulation, a 2x stock of 50 nM N-formyl-L-methionyl-L-leucyl-L-phenylalanine (fMLP; Sigma-Aldrich) was added. For F-actin inhibition, a 2x stock of 1 μM latrunculin B (Sigma-Aldrich) and 25 nM fMLP was used. All initial stocks were dissolved in 100% dry DMSO and freshly diluted in imaging media before experiments.

#### Imaging cells on nanopatterned substrates

Coverslips with nanopatterns were obtained from Cail et al., 2022. The nanopatterned coverslips were held in Attofluor™ Cell Chambers (A7816; thermofisher) and coated with 200 μL of 40 μg/μL porcine fibronectin dissolved in imaging buffer for 20 min at room temperature. The fibronectin solution was then removed and washed twice with the imaging buffer. 500 μL differentiated HL-60s (~0.8 million/mL) labeled with CellMask Deep Red (Invitrogen) were plated on to the coverslip. The chamber with the coverslip was then transferred to a 37 °C/5% CO2 incubator for 15 min to allow cells to adhere. The coverslip was then gently washed twice with imaging buffer to remove unattached cells and finally exchanged for 500 μL fresh imaging media. For the agarose compression experiment, the imaging media was removed and a precast agarose pad with 1 μM latrunculin B was gently placed on top of the cells on nanopatterns right before imaging. For cell migration on nanopatterns, 20 nM fMLP was added to cells before imaging.

#### Fixation

To fix cells, medium was aspirated followed by immediately adding 2% glutaraldehyde (Sigma-Aldrich) and incubation at room temperature for 10 min. Cells were washed twice with PBS and then quenched with 0.1% sodium borohydride (Sigma-Aldrich) for 7 min followed by two or three 10-min PBS washes. All dilutions were prepared fresh in cytoskeleton buffer with sucrose (CBS), a T.J. Mitchison laboratory (Harvard Medical School, Boston, MA) recipe: 10 mMMES, pH 6.1, 138 mM KCl, 3 mM MgCl2, and 2 mM EGTA, with 0.32 M sucrose added fresh before use.

#### Cell compression with agarose pad

A solution of 4% low-melt agarose (A-204; Gold Biotechnology) was made in imaging media and microwaved in a loosely capped conical tube placed in a water reservoir. Heating was done in short increments to promote melting while preventing the solution from boiling over. Once completely melted, the gel was kept at 4 °C to allow cooling while preventing solidification. 1 μM of latrunculin B was then added to the gel solution after cooling. The mold for gel casting was made by cutting the back end (that goes into a pipette) of a 1000 μL pipette tip to get a cylinder of ~0.7 cm tall. The mold was then placed in a 35 mm dish with the uncut edge facing the bottom. ~700 μL of gel solution was added to each mold and left at RT for solidification for 10–15 min. The mold was then removed carefully to let the gel keep solidifying for another 30 min at RT. To compress the cells, the imaging solution was removed, and the agarose pad was handled with a tweezer and placed directly on top of the cells.

#### Microscopy

All TIRF experiments (except Figure 3 F and G) were acquired at 37 °C/5% CO2 with the DeltaVision OMX SR microscope (GE Healthcare) with a 60×/1.42 NA oil Plan Apochromat objective (Olympus) and 1.518 refractive index oil (Cargille). Images were acquired with Acquire SR software and processed with soft-WoRx. TIRF-SIM images were reconstructed using OMX SI Reconstruction with the default parameters. TIRM imaging was performed using the ring TIRF light path on the DeltaVision OMX SR. FRAP experiments were performed using the ring TIRF light path combined with the FRAP/Photoactivation module on the DeltaVision OMX SR.

The single particle tracking experiments (**Figure 3 F and G**) were performed on a Nikon Eclipse Ti microscope equipped with a Borealis beam-condition unit (Andor Technology), a 100× Plan Apochromat TIRF 1.49 NA objectives (Nikon), and an iXon Ultra EMCCD camera. Environmental control (37 °C/5% CO2; Okolab) was used. Acquisition was controlled with Micro-Manager.

### Image analysis

All analysis code and CSV files of all data are available at https://github.com/MuziyueWu/WAVE.

### Quantification of the linear and planar patterns

WAVE complex signal was segmented based on the fluorescence intensity from the eGFP-Sra1 channel. For quantification of the WAVE complex at the tips of lamellipodia, segmented regions at the desired location were manually selected and quantified with Python package scikit-image (Van Der Walt et al., 2014). For identification of the WAVE complex nanorings, each segmented region was fit to a circle and filtered based on the fitting score (Supplementary Figure S5), followed by quantification as described above. For the membrane-bounded droplets, the images were segmented based on the fluorescence intensity from the Fyn-Hotag3 channel. The radius was calculated by taking the average of the long and the short axes of the segmented region.

### Single Molecular Tracking

Tracking of the single molecules was performed in Fiji using the plug-in TrackMate (Tinevez et al., 2017). Tracks were filtered for desired properties (duration, quality, etc.), and coordinates were exported as CSV files. Kymographs were created by x-axis projection of the roi. The diffusion coefficient of the molecules was calculated as described in Mehidi et al., 2021. Briefly, the Mean Square Displacement of each subunit was calculated by

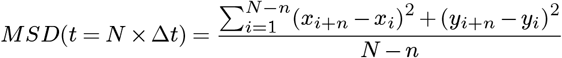

where *x*_*i*_ and *y*_*i*_ are the coordinates of the subunit position at time *i* × Δ*t*. The diffusion coefficient D is defined as the slope of the affine regression line fitted to the *n* = 1 to 4 values of the *MSD*(*n* × Δ*t*).

### Quantification of the standard fluorescent candles

Fixed cells were imaged five consecutive times and bleach-corrected by fitting into a decaying exponential curve. The fluorescent candles were then segmented based on multi-otsu thresholding. Each fluorescent spot was fit with a 2D Gaussian, and the background-independent signal was calculated using the equation for volume under the fitted Gaussian curve. The spots were then filtered based on the fitting quality and the amplitude of the Gaussian curve to remove the spots that could contain more than one candle. Finally, the spots were tracked using Python package Trackpy, and the mean fluorescent intensities of the spots that can be detected for more than three frames were used for histogram plotting.

### FRAP analysis

For the recovery pattern analysis (**Figure 4F and G**), a 1 × 30 pixel region along the middle region of the bleached array was extracted for each cell in Fiji and the fluorescent intensities at each time point were then averaged between all the cells. For the recovery kinetics analysis (**Figure 4 H and Supplementary Figure S2 E**), the normalized fluorescent intensity of the bleached region is determined using the mean fluorescent intensity of the top 10 pixels within the bleached region, normalized by the average fluorescent intensity (top 10 pixels) of a neighboring unbleached region at the corresponding time point. The halftime and amplitude of WAVE complex recovery were determined by plotting the normalized fluorescence within the bleached region as a function of time and fitting the data with an exponential function using the Stowers Institute ImageJ Plugins.

### Quantification of WAVE complex on beads

The beads under the cells were identified by finding the overlap between the beads and the cells. Beads were segmented based on the bead channel. And cells were segmented based on the eGFP-Sra1 channel. Tracking of the beads under the cells was performed using the Python package Trackpy. The fluorescent intensity of the WAVE complex/membrane dye on each bead was normalized by the WAVE signal/membrane dye surrounding that bead (not on the bead). The data used for plotting were filtered and aligned based on the peak WAVE complex signal. For tracking the fluorescent intensity of the WAVE complex (eGFP-Sra1 channel) on beads over time, the normalized WAVE complex and membrane dye signal on each bead was plotted as a function of time. For comparison between the WAVE complex and membrane signal, the average of three top fluorescent signals for each channel was calculated and then averaged for each cell.

### Quantification of WAVE complex on nanopillars

Linescans of the membrane dye and the WAVE complex (eGFP-Sra1) channel were generated by projection of the fluorescent signal onto the long axis of each nanopillar, followed by 1D Gaussian smoothing of the linescan curve.

## Acknowledgement

We thank the Woolfson lab and the Weiner lab for helpful discussion. We thank Kirstin Meyer, Sue Sim, and Evelyn Strickland for a critical reading of the manuscript. We thank the Drubin lab and the Shu lab for providing the plasmids. We thank Kari Herrington and SoYeon Kim of the UCSF Imaging Core for their microscopy expertise. This work was supported by an American Heart Association Predoctoral Fellowship 23PRE1018810 (M WU.), the National Institute of General Medical Sciences GM118167 (O.D. Weiner), National Science Foundation/Biotechnology and Biological Sciences Research Council grant 2019598 (O.D. Weiner), and the National Science Foundation Center for Cellular Construction (DBI-1548297). The work was also funded by a 19-BBSRC-NSF/Bio joint grant to Weiner and Woolfson: Biotechnology and Biological Sciences Research Council grant BB/V004220/1 (D.N. Woolfson); National Science Foundation grant 2019598 (O.D. Weiner). Data for this study were acquired at the Center for Advanced Light Microscopy at UCSF on an OMX-SR obtained using grants from the NIH (5R35GM118119), the UCSF Program for Breakthrough Biomedical Research funded in part by the Sandler Foundation, the UCSF Research Resource Fund Award, and HHMI. The funders had no role in study design, data collection and analysis, decision to publish, or preparation of the manuscript.

## Supplementary Figures

**Figure S1.**
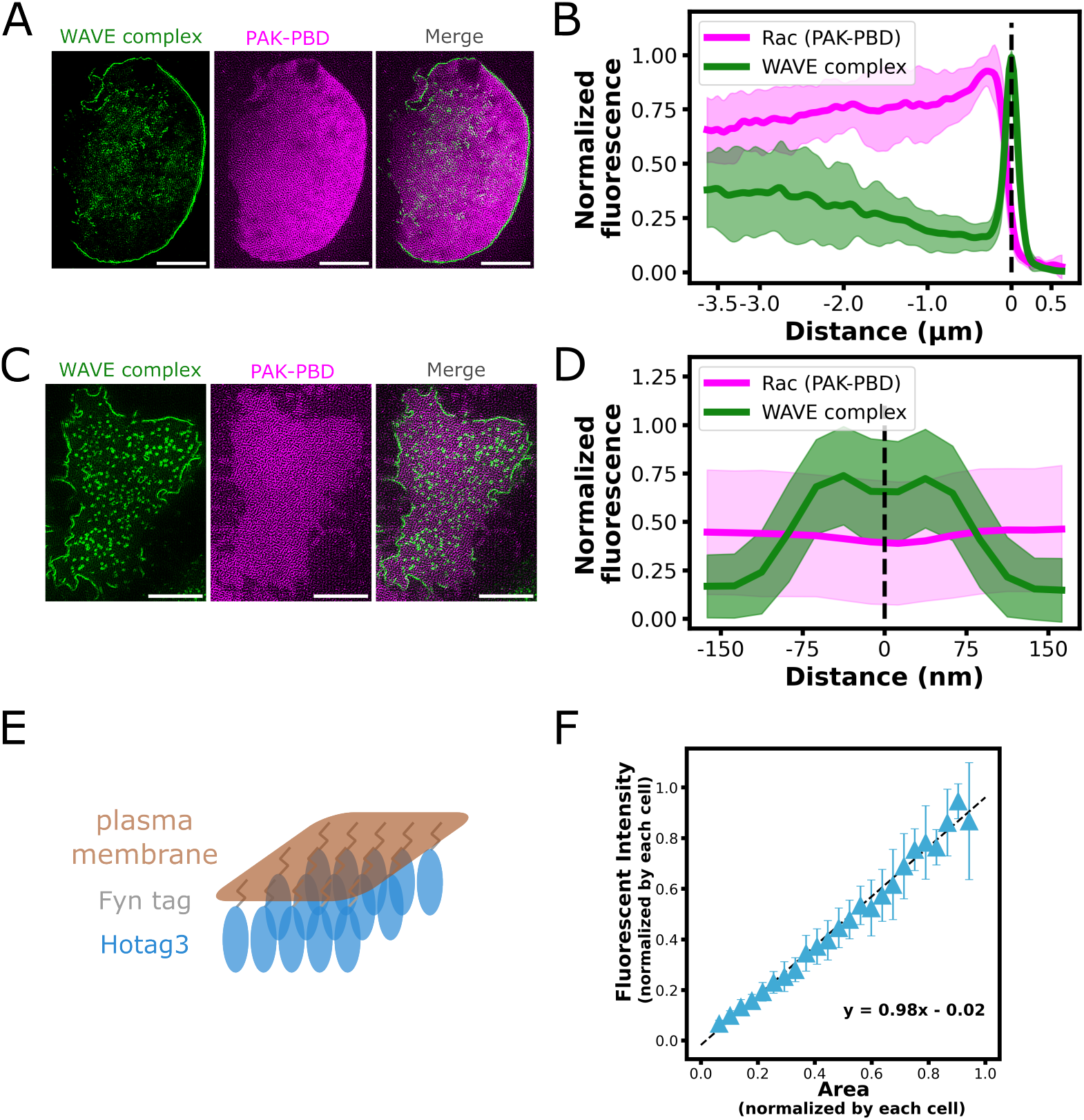
The WAVE complex adopts a restricted linear pattern compared to Rac and other planar structures. (**A**) The WAVE complex specifically localizes to the tips of lamellipodia, while active Rac (PAK-PBD) spans a permissive region at the front of a migratory HL-60 cell. TIRF-SIM imaging; scale bar: 5 μm. (**B**) Linescan across the front half of the cells. Rac shows a gradual signal increase towards the front side of the cell while the WAVE complex shows a sharp signal increase at the tips of lamellipodia. N= 17 cells. (**C**) The WAVE complex forms nanoring structures in HL-60 cells treated with latB while Rac (PAK-PBD) spans uniformly on cell membranes. TIRF-SIM imaging; scale bar: 5 μm. (**D**) Linescan across the middle section of the WAVE complex nanorings. WAVE complex signal shows ring patterns while Rac is uniformly distributed. N = 1700 nanorings from 12 cells. (**E**) Schematic showing how Fyn-Hotag3 forms planar structures on cell membranes. (**F**) Fyn-Hotag3 exhibits a linear relation between the fluorescent intensity and the area, but not the length of the pattern.

**Figure S2.**
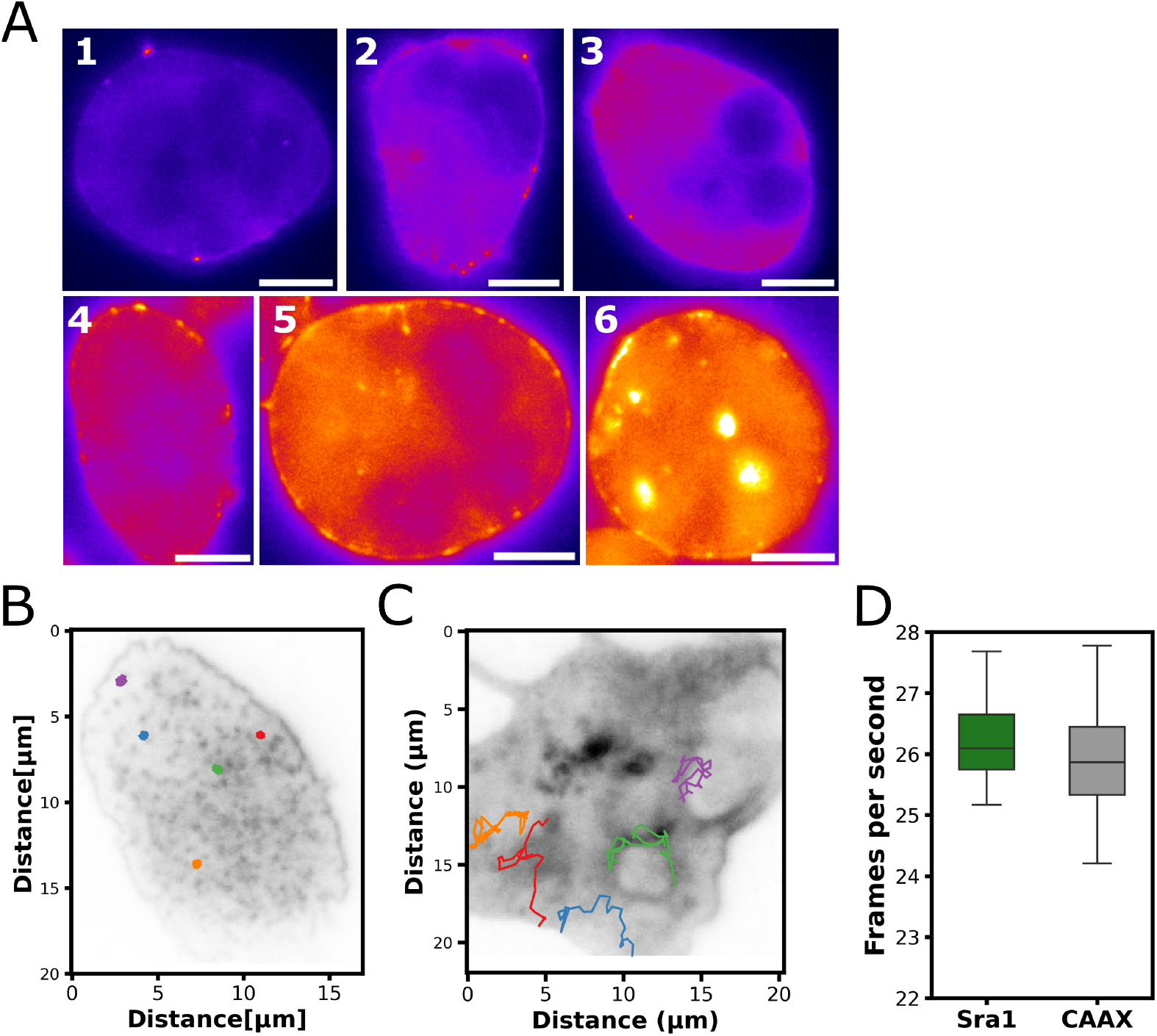
Supplement Hem1 KO and single molecule tracking data. (**A**)Epi-fluorescence imaging data showing the various expression levels from low to high of selected cells in Figure 3 C and D. Scale bar: 5 μm. (**B**) Selected tracks of single Halo-Sra1 proteins in the background of fully labeled Halo-CAAX signal. (**C**) Selected tracks of single Halo-CAAX proteins in the background of fully labeled Halo-CAAX signal. (**D**) Image acquisition speed for the single-molecule tracking experiment. N = 11 for Halo-Sra1 cells, n = 24 for Halo-CAAX cells.

**Figure S3.**
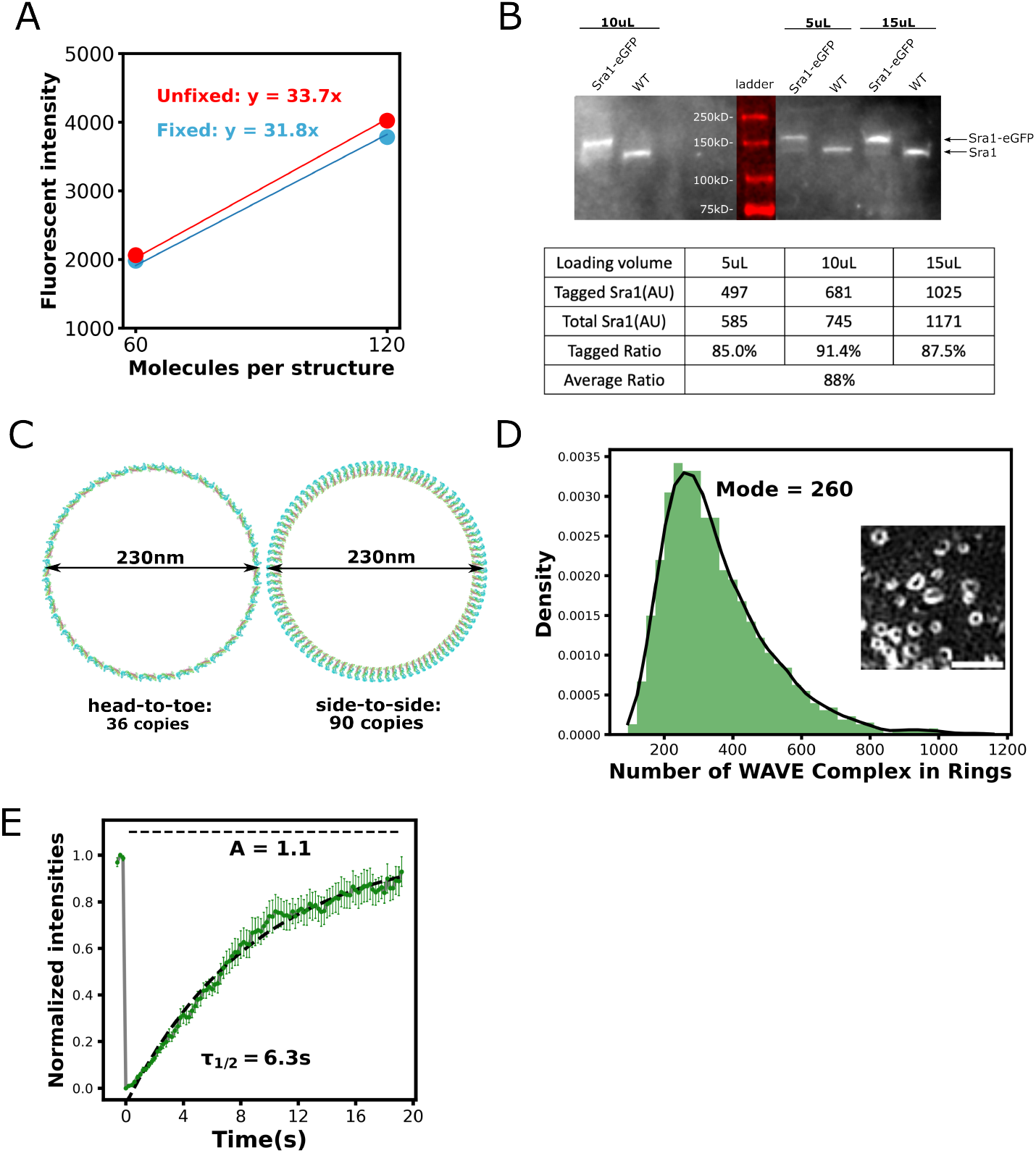
Supplement molecular counting and FRAP data. (**A**) Correction for fluorescence loss after fixation. 60mer: n = 73 puncta from n = 6 fixed cells and n = 88 puncta from n = 8 unfixed cells. 120mer: n = 35 puncta from n = 7 fixed cells and n = 40 puncta from n = 6 unfixed cells. Ring-TIRF imaging. (**B**) Western blot with serial dilutions of the sample to calculate the percentage of eGFP-tagged Sra1 subunits in cells. The percentage of eGFP-tagged Sra1 subunits in the eGFP-Sra1 cell line used for molecular counting is 88%. (**C**) The numbers of the WAVE complex required to make a single-layered ring of 230nm in diameter, based on PDB 3p8c by Chen, Z., et al. (2010). (**D**) Histogram distribution of the copies of WAVE complexes in nanorings, with a mode of 260. n = 5217 from 38 cells acquired in four experiments. Black line shows the kernel density estimation of the distribution. Middle right inset shows the TIRF-SIM live imaging of the eGFP-Sra1 labeled WAVE complex nanorings in cells treated with latB (scale bar: 1 μm). (**E**) The WAVE complex at the tips of lamellipodia in migratory HL-60 cells shows a full fluorescent recovery after photobleaching. N = 15 cells.

**Figure S4.**
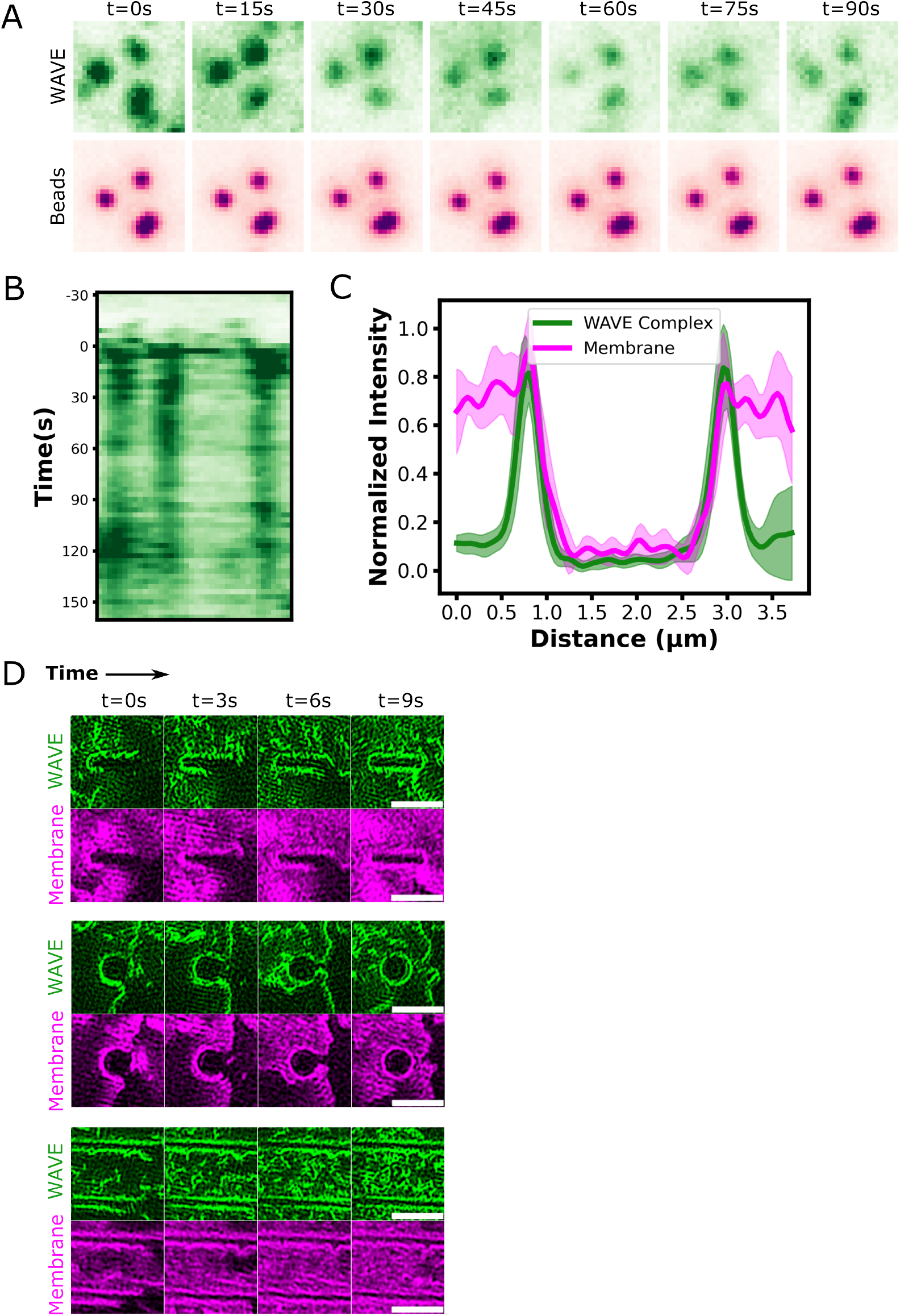
WAVE Complex associates with negative curvatures induced by beads and nanopatterns. (**A**) The WAVE complex (eGFP-Sra1; green) can stably associate with 200nm beads (red) over 90s. (**B**) Kymograph of the WAVE complex on beads, again showing persistent recruitment of the WAVE complex on beads. (**C**) Linescan of WAVE complex and membrane dye on rectangular nanopillars. The WAVE complex shows a sharp signal increase at the boundary where the plasma membranes exit the TIRF plane, corresponding to sites of negative membrane curvature induced by the nanopillars. N = 8 nanopillars. (**D**) The WAVE complex and the membrane dye on different nanopatterns in migratory HL-60 cells over time. In migratory HL-60 cells, the WAVE complex can also associate with negative curvature sites induced by nanopatterns. TIRF-SIM imaging; scale bar: 2 μm.

**Figure S5.**
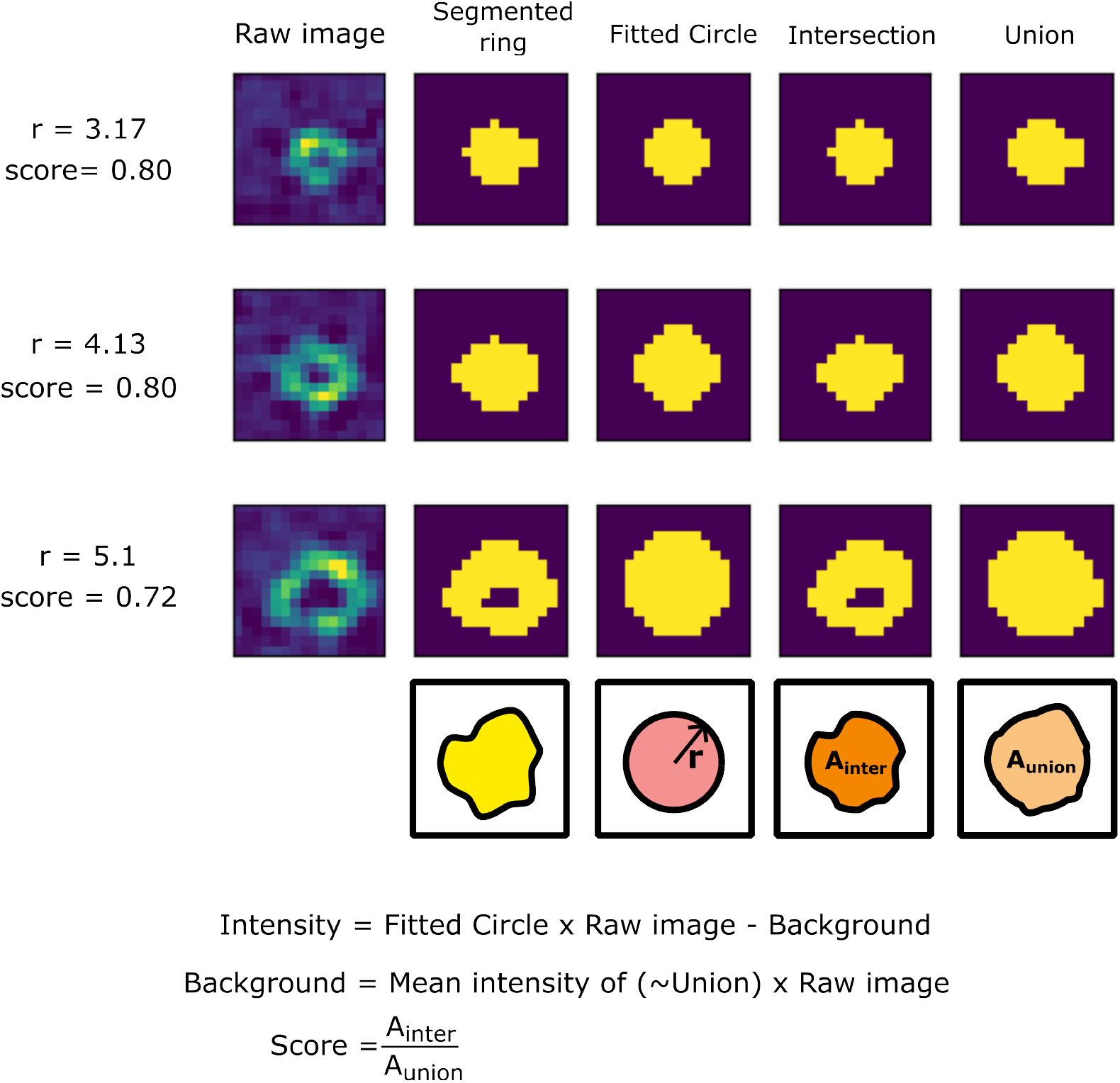
Identifying WAVE complex nanoring structures by circle fitting. The raw images were first segmented based on a certain threshold. Each segmented region was then fitted to a circle with a radius optimized to achieve the highest filtering score, defined as the ratio between the intersection and union of the segmentation and the fitted circle. The segmentations above a certain score were used for quantification. The fluorescent intensity of each nanoring is defined as the sum of the intensity values within the fitted circle minus the background.

## Supplementary Movies

**Video 1. WAVE complex organization pattern in HL-60 cells**. Left shows the eGFP-labeled Sra1 subunits in a migratory HL-60 cell treated with chemoattractant (25 nM fMLP), where the WAVE complex forms a linear array at the tips of lamellipodia. Right shows the eGFP-labled Sra1 subunits in an HL-60 cell treated with actin inhibitor (0.5 μM latB), where the WAVE complex forms nanoscale rings at membrane invagination sites. TIRF-SIM imaging; scale bar: 2 μm.

**Video 2. Single molecule tracking on sparse-labeled Halo-tagged cell lines**. Single molecular tracking data of the Halo-Sra1 cells (left) and Halo-CAAX cells (right) sparse-labeled with JF549 and treated with 0.5 μM latB. Magenta circles annotate the single molecules identified by TrackMate and cyan lines show selected tracks. TIRF imaging; scale bar: 2 μm.

**Video 3. FRAP on WAVE complex at the tips of lamellipodia in a migratory HL-60 cell**. Photobleaching happened at t = 0.6 s. White circle annotates the region for photobleaching. Images were acquired at 200 ms/frame. Ring-TIRF imaging; scale bar: 5 μm.

**Video 4. FRAP on WAVE complex in a latB-treated HL-60 cell**. Photobleaching happened at t = 2.0 s. White box annotates the region for photobleaching. Images were acquired at 500 ms/frame. Ring-TIRF imaging; scale bar: 5 μm.

**Video 5. TIRF imaging of an HL-60 cell migrating on 200 nm beads**. WAVE complex (eGFP-Sra1; green) colocalizes with and stays on 200nm beads (magenta) as cells migrate. Ring-TIRF imaging; scale bar: 5 μm.

**Video 6. TIRF-SIM imaging of an HL-60 cell migrating on 200 nm beads**. WAVE complex (eGFP-Sra1; green) forms ring structures around 200nm beads (magenta) as cells migrate. TIRF-SIM imaging; scale bar: 5 μm.

**Video 7. HL-60 cells migrating on nanopatterns**. WAVE complex enriches at the rim of nanopillars (left) and nanoridges (right) as cells migrate. TIRF-SIM imaging; scale bar: 5 μm.

